# Pure wild forest reindeer (*Rangifer tarandus fennicus*) or hybrids? A whole-genome sequencing approach to solve the taxonomical status

**DOI:** 10.1101/2023.08.16.553517

**Authors:** Melak Weldenegodguad, Milla Niemi, Sakari Mykrä-Pohja, Kisun Pokharel, Tuula-Marjatta Hamama, Antti Paasivaara, Juha Kantanen

**Author notes:** Corresponding authors: Milla Niemi,; Juha Kantanen.

## Abstract

In Finland, the geographic distribution of semi-domestic reindeer (*Rangifer tarandus tarandus*) and wild forest reindeer (*Rangifer tarandus fennicus*) partly overlap in the vicinity of the southern border of the reindeer herding area. Hybridisation of these two subspecies is one of the major threats for the wild forest reindeer population, and the implemented conservation program targets preventing the crossbreeding of these reindeer. In this paper, we resequenced genomes of four *Rangifer tarandus* individuals with unknown taxonomical status and investigated their ancestries by comparing the genomic data with the existing resequenced data of the Finnish semi-domestic reindeer and Finnish wild forest reindeer. The genetic relationship investigations suggest that the four individuals were pure forest reindeer animals. Our study provides critical knowledge for practical conservation actions of wild forest reindeer in the reintroduction project, for which it is essential to recognise each individual’s origin. In the future, it will also offer novel insights into the spread of native wild forest reindeer to new geographic regions in Finland. For subsequent studies, additional resequenced genomic data of *Rangifer* individuals will be needed to develop an ancestry information marker panel of single nucleotide polymorphisms for rapid and cost-effective identification of hybrid individuals of semi-domestic reindeer and wild forest reindeer.

## Introduction

Hybridisation or crossbreeding between individuals of different species may occur naturally in 10% of animal species (Mallet 2005). While it could act as a source of genetic variation, especially hybridisation between wild populations and their domesticated or captive-reproduced relatives is an increasing conservation concern worldwide (e.g., Randi 2008). The introduction of non-native species to new areas has led to hybridisation or crossbreeding, as with several European ungulate populations, for example (e.g., Randi 2005; Iacolina et al. 2019).

The forest-dwelling wild forest reindeer *Rangifer tarandus fennicus*, a native subspecies of reindeer with circumpolar distribution, nowadays exists only in Finland, Russian Karelia and the westernmost part of the Arkhangelsk oblast (Panchenko et al. 2021). In the winter of 2021, there were about 2,900 individuals in Finland, divided into two main subpopulations, Kainuu and Suomenselkä (Paasivaara et al. 2021, 2022; Pöllänen et al. 2023). The Suomenselkä population was introduced from Kainuu in the early 1980s. In addition, two new subpopulations have been reintroduced recently (Mykrä-Pohja and Niemi 2022). Based on aerial censuses conducted in late winter 2023, the current total number of Finnish wild forest reindeer is about 3,000 individuals (Natural Resources Institute Finland, unpublished data), divided over the two main subpopulations (Paasivaara et al. 2021). The Russian population has decreased during the last few decades, the last estimate being approximately 2,300 individuals (Panchenko et al. 2017; Danilov et al. 2020), leading to the global population of about 5,000–6,000 individuals. In contrast to decreasing wild reindeer populations, Klokov (2007) estimated that there are nearly 2,000,000 semi-domestic reindeer globally, of which one-third are distributed in Fennoscandia.

Although the Fennoscandian semi-domestic reindeer has been domesticated from the tundra reindeer (also called the mountain reindeer, *Rangifer tarandus tarandus*) (Røed et al. 2021), the semi-domestic reindeer and the wild forest reindeer partly have a common history as suggested by Heino et al. (2021). They studied ancient reindeer bone samples from indigenous Sámi offering sites and found that mitochondrial DNA (mtDNA) haplotypes common to wild forest reindeer were present even in the most northern part of present-day Finland, that is, in the current reindeer husbandry area. They began to be replaced by mtDNA haplotypes found in semi-domestic reindeer starting between 1400 and 1600 CE, indicating the growing importance of reindeer herding during those centuries. Correspondingly, the wild forest reindeer were hunted in a reindeer herding area, and the last herd was not shot until 1883 (Montonen 1974).

The recent whole-genome sequencing study evidenced that the wild forest reindeer are genetically distinct from the semi-domestic reindeer (Pokharel et al. 2023). Moreover, the archaeo-osteological study by Rankama and Ukkonen (2001) indicated separate origins for northern Fennoscandian forest and tundra reindeer; the forest reindeer may have originally spread to Finland from the east circa 7,500 years ago, following the retracting ice margin, while the tundra reindeer may have colonised northern Fennoscandia especially along the Norwegian narrow ice-free coastal zone.

Crossbreeding of semi-domesticated reindeer and wild forest reindeer has been recognised as a major threat for the wild forest reindeer genome (Härkönen and Bisi 2007), and an active prevention program targets preventing hybridisation and removing possible hybrids from the Finnish subpopulations of wild forest reindeer (Niemi et al. 2021). Most encounters leading to crossbreeding between the semi-domestic reindeer and wild forest reindeer occur near the southern border of the reindeer husbandry area. However, there is sporadic, small-scale reindeer keeping also in the southern part of Finland, meaning that hybridisation could occur there as well if animals escape from their owner (Wildlife Service Finland, unpublished data). Even if the appearance of hybrids in the wild forest reindeer population seems to be low (Pokharel et al. 2023), the expanding distribution area of wild forest reindeer will most likely lead to an increased number of encounters with semi-domesticated and wild reindeer, underlining the need for tools to test a possible hybridisation.

In this study, we analysed the genome of four supposed wild forest reindeer individuals to foreclose their possible hybridisation status and to recognise where a possible subpopulation might originate. Three of these animals were intended to be used as founder individuals in a wild forest reindeer reintroduction project, underlining the importance of their genetic origin. For determination of their taxonomical status (and possible hybridisation), we used the existing whole-genome sequencing data of the Finnish semi-domestic reindeer and wild forest reindeer (Weldenegodguad et al. 2020; Pokharel et al. 2023).

## Materials and Methods

### Collecting samples

In the winter of 2019, a wild forest reindeer female and its female calf (*individual 1*; see Table 1 and Supplementary Tables S1 and S2) were detected in the reindeer husbandry area in Kainuu, where a 90-kilometre-long fence has been built to separate semi-domestic reindeer and wild forest reindeer populations (Niemi et al. 2021). Both individuals were caught and transported (Permit criteria of the Finnish National Animal Experiment Board; Decisions on the granting of license of team Regional State Administrative Agency for Southern Finland, ESAVI: 6336/04.10.03/2012, 587/04.10.07/2016 and 23666/2018) to the breeding enclosure located in Lauhanvuori National Park (62.1521° N, 22.1751° E), which is one of two reintroduction sites of wild forest reindeer in the EU-funded project WildForestReindeerLIFE (LIFE15 NAT/FI/000881) (Mykrä-Pohja and Niemi 2022). Because those transported individuals were intended to act as founder individuals for the newly reintroduced wild forest reindeer population and animals were found from the reindeer husbandry area, the calf’s possible hybridisation status needed to be tested. As the adult female was pregnant and gave a birth in the enclosure in the spring of 2019, the status of that newborn male calf (*individual 2*) also had to be checked. The young female was anaesthetised and sampled in Autumn 2020 and the male in the summer of 2021.

**Table 1.** Summary of samples used in whole-genome sequence analysis.

| Population | No. of individuals |
| --- | --- |
| Finnish semi-domestic reindeer (*R. t. tarandus*) | 10 |
| Finnish Forest reindeer (*R. t. fennicus*) | 6 |
| Studied *Rangifer* individuals | 4 |
| Alaskan wild caribou (*R. t. caribou*) | 1 |

In January 2021, the police captured a young *Rangifer* male (*individual 3*) from Helsinki (https://yle.fi/a/3-12262918), which is located more than 200 kilometres to the south of the wild forest reindeer reintroduction areas and 300 kilometres from the distribution area of the closest wild population (Fig. 1) (Pöllänen et al. 2023). After recovering in the Helsinki Zoo, that individual was also transported to the wild forest reindeer breeding enclosure in Lauhanvuori National Park. Its mtDNA was analysed before transportation (unpublished data, Korkeasaari Zoo Korkeasaari Zoo 2022: https://www.korkeasaari.fi/helsinkiin-eksynyt-orpo-vasa-varmistui-metsapeuraksi/), but the analysis did not reveal possible crossbreeding with a wild forest female and semi-domestic reindeer male (Amorim et al. 2020); the possible hybridisation status needed to be confirmed through this study.

**Fig. 1.**
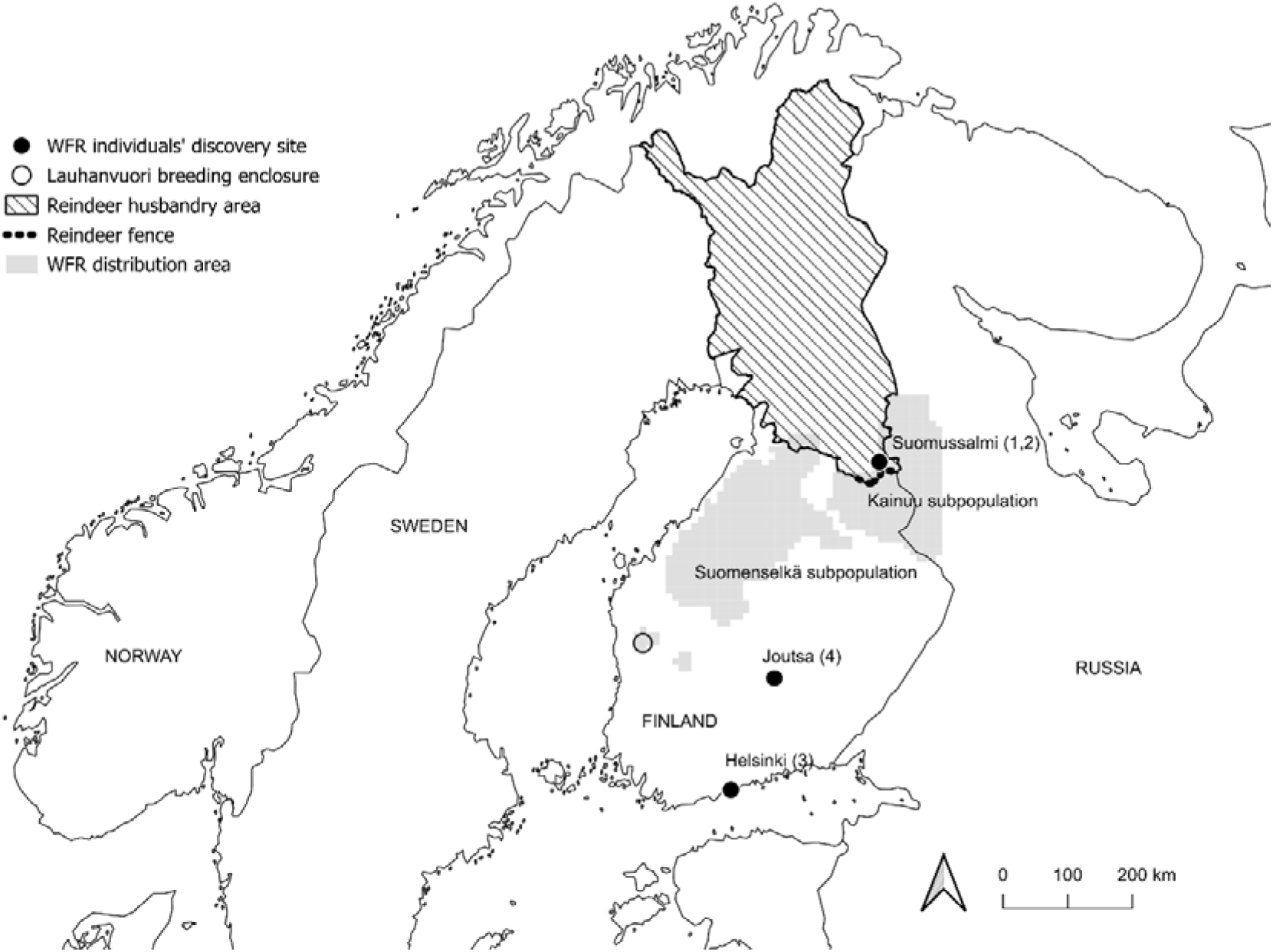
Discovery sites of studied wild forest reindeer individuals and breeding enclosure in Lauhanvuori National Park. The subpopulations of wild forest reindeer are marked with grey and reindeer husbandry areas with diagonal lines. This information was provided by the Natural Resources Institute Finland (wild forest reindeer distribution), the Reindeer Herders’ Association (reindeer husbandry area) and Eurostat (country borders). The abbreviation ‘WFR’ refers to ‘wild forest reindeer’.

With these three samples related to the reintroduction of wild forest reindeer, an older sample (*individual 4*) from a supposed wild forest reindeer was also analysed. This individual was found dead in Joutsa, Finland, in the autumn of 2017 outside the known distribution area of wild forest reindeer (Fig. 1), and it was necessary to identify its origin.

### Whole-genome sequencing

To examine the genetic relationships between the Finnish semi-domestic and the four wild forest reindeer individuals described above, we used a total of 21 resequenced reindeer samples (Table 1 and Supplementary Table S1). The previous data included genome sequences of 10 Finnish semi-domestic reindeer, six wild Finnish forest reindeer and, as an outgroup, one wild caribou from Alaska, USA (Weldenegodguad et al. 2020; Pokharel et al. 2023); these data were generated at Beijing Genomics Institute using the Illumina HiSeq 4000 platform.

The new samples described above and investigated here were sequenced using the Illumina HiSeq 4000 with a 150 bp paired-end strategy at Novogene.

### Bioinformatic analysis

Prior to the bioinformatic analysis, the quality of the raw reads was assessed using FastQC software v0.11.8 (Andrews 2010). MultiQC v1.9 (Ewels et al. 2016) was used to summarise the quality control reports of all samples from FastQC. High-quality clean reads for each sample were mapped against the reindeer reference genome (Weldenegodguad et al. 2020) using BWA v0.7.17 (Li and Durbin 2009) with default parameters. Following mapping, the SAM files generated by BWA were converted to binary equivalent BAM files and sorted using Picard tools v2.21.4 (https://broadinstitute.github.io/picard/). PCR duplicates from the aligned reads were removed using Picard tools.

After preprocessing the mapped reads, the Genome Analysis Toolkit (GATK) best practices pipeline (Van der Auwera et al. 2013) was used to identify high-quality variants (single nucleotide polymorphisms [SNPs] and indels) from the uniquely mapped reads. Variants were called using the GATK v4.2.0.0 HaplotypeCaller from the uniquely mapped reads. Based on GATK4 user guide recommendations, sequencing and alignment artifacts from the variants were discarded using the following parameters: FS > 60.0, MQ < 40.0, MQRankSum < −8.0, QD < 2.0, ReadPosRankSum < −8.0 and SOR > 3.0 for SNPs and FS > 200.0, MQ < 40.0, QD < 2.0, ReadPosRankSum < −8.0 and SOR > 5.0 for indels.

To examine the genetic relatedness between the samples, a principal component analysis (PCA) was conducted using the identified SNPs with a Bioconductor R package SNPRelate, v1.28.0 (Zheng et al. 2012). SNPs were first filtered based on linkage disequilibrium (LD) using the snpgdsLDpruning function in the SNPRelate package to avoid strong influence of linked SNP clusters. We used 0.2 LD thresholds for filtering the SNPs. The R function snpgdsPCA was used to perform a PCA plot.

Moreover, the identified SNPs were utilised to generate a Neighbour-Joining phylogenetic tree using SNPhylo v. 20160204 (Lee et al. 2014). Before constructing the tree, SNPs were further filtered using SNPhylo with the following filter parameters: minimum depth of coverage > 5, percentage of low-coverage samples < 5%, percentage of samples with no SNP information < 5%, LD < 0.1 and minor allele frequency > 0.05. The filtered SNPs were then concatenated to generate sequences and used to perform multiple alignments using MUSCLE v3.8.31 (Edgar 2004). A Neighbour-Joining tree was then constructed by running DNAML programs in the PHYLIP package (Felsenstein 2005). Bootstrap analysis was performed using the “phangorn” package with 100 replications (Guindon et al. 2010). FigTree v1.4.4 (Rambaut 2018) was used to visualise the phylogenetic tree.

Furthermore, population admixture based on the identified SNPs was examined using ADMIXTURE software, v.1.3.0 (Alexander et al. 2009). The SNPs in VCF format were first converted into binary PLINK format for ADMIXTURE input using PLINK v1.90b6.21(Purcell et al. 2007). ADMIXTURE was then examined using different ancestral clusters (K) ranging from two to five; to estimate standard errors, the bootstrap parameter was set to 100 replicates.

## Results and Discussion

### Whole-genome sequence data

In this study, a total of 441 gigabases (Gb) of paired-end whole-genome sequence data were generated from 21 samples (Table 1 and Supplementary Table S2) to examine the genetic relatedness between the samples. The quality of all the samples was found to be high and, hence, quality filtering was not performed. On average, each sample had 274□million (M) and 41.18□Gb clean reads and bases, respectively (Supplementary Table S2). The reads were successfully aligned to the assembled reference genome, with an average alignment rate of 98.31%, and represented 10-fold coverage (Supplementary Table S2).

### Variant calling

A total of 28.1 M SNPs were detected using the uniquely mapped reads across all 21 samples (Supplementary Table S3). The average number of SNPs detected per individual was 8.1□M. In the semi-domestic reindeer, the average number of SNPs was 7.9 M, in the wild forest reindeer it was slightly higher at 8.2 M and in the four samples investigated here it was 8.5 M. On average, the calculated transition-to-transversion (Ts/Tv) ratios were 1.90 per individual (Supplementary Table S3). The observed Ts/Tv ratios were found to be slightly lower than that of humans (2.1) (Lachance et al. 2012) and bovines (2.2) (Stothard et al. 2011; Choi et al. 2014, 2015). Moreover, a total of 3.9□M indels were detected across all 21 samples, and on average 1.23 M indels were detected per individual.

### Population structure analysis

PCA and phylogenetic tree analysis were used to examine the genetic relationships between all 21 animals. In the phylogenetic tree, the wild caribou from Alaska was an outgroup. The PCA plot and phylogenetic tree revealed two major groups: the Finnish semi-domestic reindeer versus the Finnish wild forest reindeer (Fig. 2 and Fig. 3). Both the PCA plot and phylogenetic tree clearly indicated that the two animals from the Lauhanvuori breeding enclosure, the *Rangifer* sp. individuals found in Helsinki and Joutsa (the latter one was dead), were pure *Rangifer tarandus fennicus*, and no sign of hybridisation with semi-domestic reindeer was detected.

**Fig. 2.**
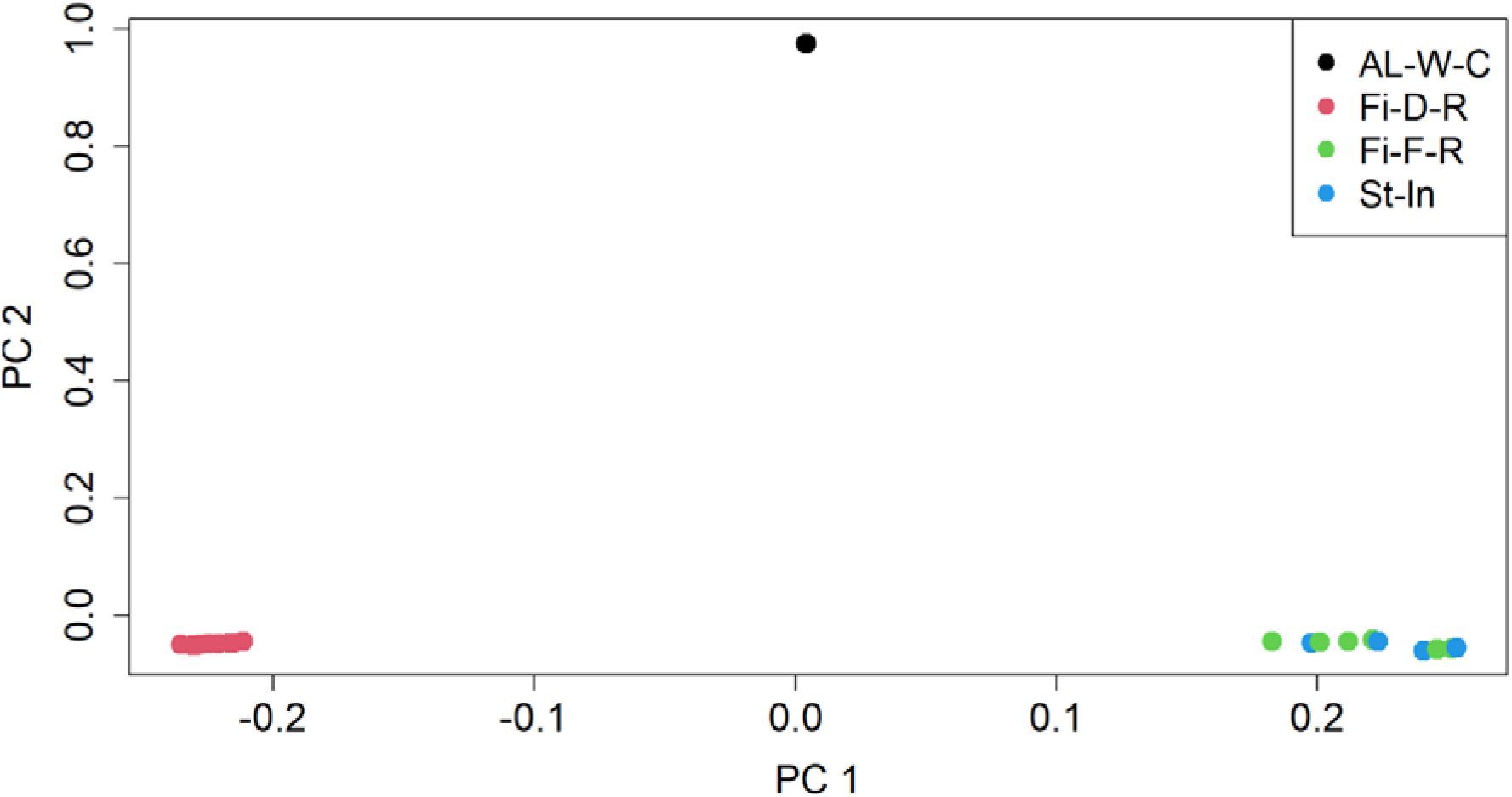
PCA plot for 21 resequenced animals. (Fi-D-R, Finnish semi-domestic reindeer; Fi-F-R, Finnish wild forest reindeer; St-In, Studied individuals; AL-W-C, Alaska wild caribou).

**Fig. 3.**
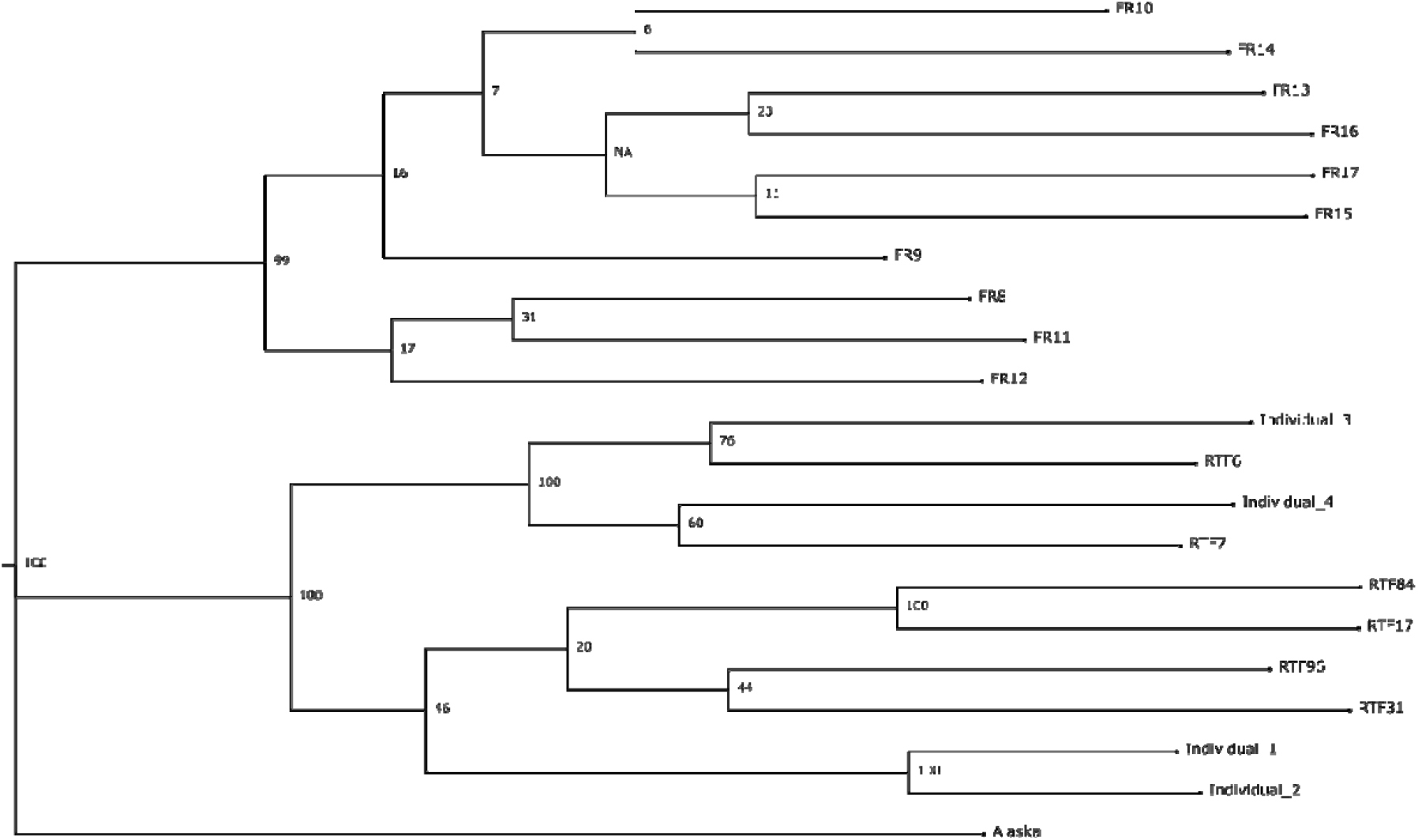
Phylogenetic analysis indicated two main clusters of reindeer populations. 1) Finnish semi-domestic reindeer (FR-individuals) and 2) Finnish wild forest reindeer (RTF-individuals)/studied individuals. Bootstrap confidence values are shown at each branch.

Population structure and admixture were further studied for all four populations (n = 21) using K values ranging from two to five. Fig. 4 shows the obtained structure plots of the four populations for all the K values. Cross-validation errors of the cluster numbers were lowest for K values two and three, increasing after that as K increased. This suggests that either two or three would be the optimal number of ancestral populations. As shown in Fig. 4, at K = 3, individuals 1, 2, 3 and 4 showed ancestries with Finnish forest reindeer populations. This clustering is consistent with the results of PCA and phylogenetic tree (Fig. 2 and Fig. 3).

**Fig. 4.**
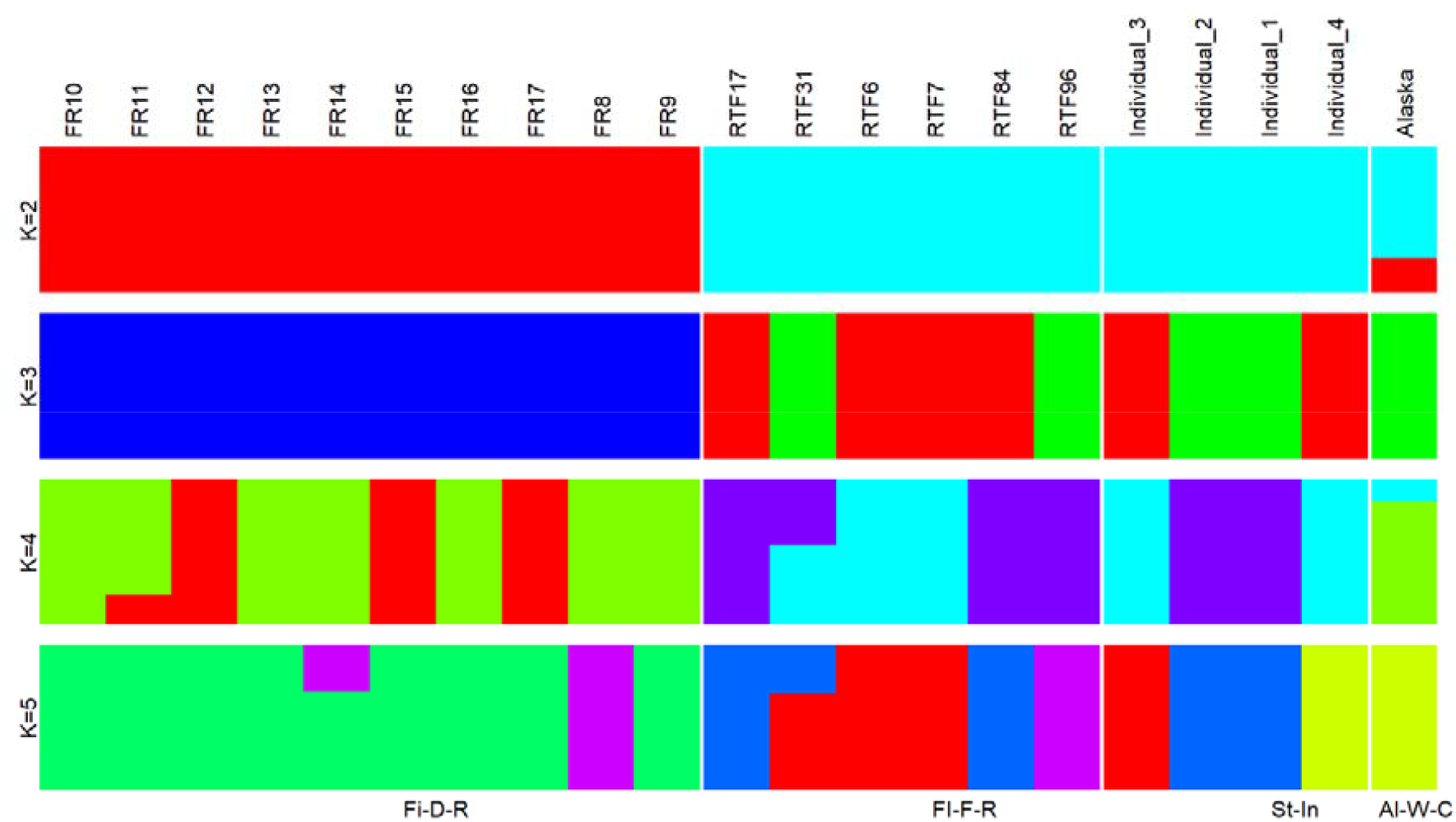
Population structure analysis of the four populations using ADMIXTURE. (Fi-D-R, Finnish semi-domestic reindeer; Fi-F-R Finnish wild forest reindeer; St-In, Studied individuals, AL-W-C, Alaska wild caribou). The analysis was repeated with different assumed numbers of clusters (K).

### Conservation implications

Our results showed that all four tested individuals were wild forest reindeer (*R. t. fennicus*), and these animals also displayed high levels of genetic variation. No marks of hybridisation were detected. Thus, three wild forest reindeer, which were kept in the breeding enclosure in Lauhanvuori National Park, were considered to be suitable as founder individuals when establishing a new, reintroduced subpopulation. Two tested males were then released into the national park, but unfortunately, the female died in the enclosure.

Our results indicate that the tested female (*individual 1*) and the young male (*individual 2*) which were born to the adult female captured from the reindeer herding area, originated from the Kainuu subpopulation, where about one-third of the population breeds in Russian Karelia near the Finnish–Russian border (the original Kuhmo–Lake Kiitehenjärvi population; see Heikura 1997). The result was highly expected as the adult female had been previously GPS-collared in Sotkamo 2011 at the age of three, and its movement history in the range of the Kainuu population was known from 2011 to 2015 (female 36/rt_fi_11_002, unpublished data base, Luke) until the battery was depleted and it was recaptured and moved to Lauhanvuori in the winter of 2019. Females usually bred in Kalevala National Park of Russian Karelia and migrated annually to Kuhmo and Sotkamo in Finland, spending winter together with large flocks of other wild reindeer (Pöllänen et al. 2023). Before being recaptured in Suomussalmi in the winter of 2019, the adult female (the dam of the *individual 1* and *2*) probably moved through the reindeer fence along the Finnish–Russian border from Russian Karelia to the reindeer management area. However, it is usual that some domestic reindeer escape from Finland to Russia through the reindeer fence every year, because domestic reindeer are observed in migrating and wintering wild reindeer flocks in Finland during yearly population monitoring seasons (Luke and National Park Services, unpublished data). Therefore, there is a high risk of crossbreeding in Russian Karelia, too.

In contrast, the young male (*individual 3*) captured from Helsinki and the one (*individual 4*) that was found after its death in Joutsa originated from the introduced Suomenselkä subpopulation. Both individuals were far away (about 200 kilometres) from the known areas of this subpopulation. The origin area of *individual 3* was unknown and somewhat confusing because calves under the age of one usually follow their mothers or the natal herd before first breeding at age two to four and after natal dispersal (Montonen 1974). Therefore, according to the known movements and young age of *individual 3*, the indications are that it was lost from the natal or normal wintering area accidentally. The poor condition of this calf at catching supports the previous explanation. Instead, *individual 4* was probably a dispersing young male (at the age of two to three years) from the Suomenselkä or Soini-Ähtäri-Karstula areas. However, one should consider the previous explanation with caution, because natal dispersal patterns of wild reindeer are still mainly unknown (van Oort et al. 2011), and adult females usually show a high rate of breeding-site fidelity (Montonen 1974; Schaefer et al. 2000).

However, there are several observations of young individuals (males and females) outside the known distribution of Suomenselkä, where the population has expanded its range during several decades (unpublished data, Luke; see also distribution maps in Härkönen and Bisi 2007 and Pöllänen et al. 2023). The findings of this study, together with the expanding range of the Suomenselkä population, indicate that the rate of true dispersal of wild forest reindeer may exist, and potentially, the probability of gene flow may increase accordingly. However, more marking and monitoring is needed to find the patterns of dispersal and gene flow for wild forest reindeer.

In this study, we have used the whole-genome sequencing approach for the detection of possible hybrid origins of individuals that are important for conservation and for investigation of the spread of wild forest reindeer to new geographic regions. Our present data for identification of the taxonomical status of the individuals with unknown origins came from whole-genome sequencing of the Finnish semi-domestic reindeer and wild forest reindeer. However, there is a need to develop an ancestry information marker (AIM) SNP panel with a limited number of SNPs for the identification of semi-domestic reindeer and wild forest reindeer hybrids (see, e.g., Somenzi et al. 2020). The AIM SNP panel can also be used to reveal the poaching of wild forest reindeer in the case that only a skinned carcass or meat is found. For the development of the AIM SNP panel, more genomes of semi-domestic reindeer and wild forest reindeer individuals need to be sequenced. Genome sequence data of hybrid animals with known pedigree records will also be useful when testing the capacity of the AIM SNP panel to identify hybrid animals.

## Supplementary information

The online version contains supplementary material available at

## Acknowledgements

Our special thanks are going to veterinarian Heini Niinimäki, who sit several days and nights with MN in Lauhanvuori breeding enclosure, waiting the target animals to come close enough to put them to sleep and collect blood samples. In addition, we want to thank Seppo Salonen and Paula Salonen from Joutsa who sent a hair sample from reindeer carcass. Mervi Honkatukia, Jaana Peippo, Tiina Reilas and Knut Roed are acknowledged for collaboration to collect the semi-domestic reindeer and caribou samples. The authors wish to acknowledge the CSC-IT Center for Science, Finland, for computational resources.

## Author contributions

M.N., S.M-P. and J.K. conceived and designed the project. M.N., S.M-P. and A.P. collected the samples of the studied four individuals. T.H. performed the laboratory analyses. M.W. did the bioinformatic analyses. M.W., M.N., S.M-P., A.P., K.P. and J.K. wrote the manuscript. All authors reviewed and approved the final manuscript.

## REFERENCES

Alexander DH, Novembre J, Lange K (2009) Fast model-based estimation of ancestry in unrelated individuals. Genome Res 19:1655–1664. https://doi.org/10.1101/gr.094052.109

Amorim A, Pereira F, Alves C, García O (2020) Species assignment in forensics and the challenge of hybrids. Forensic Sci Int Genet 48:102333. https://doi.org/10.1016/j.fsigen.2020.102333

Andrews S (2010) FASTQC. A quality control tool for high throughput sequence data

Choi J-W, Choi B-H, Lee S-H, et al (2015) Whole-Genome Resequencing Analysis of Hanwoo and Yanbian Cattle to Identify Genome-Wide SNPs and Signatures of Selection. Mol Cells 38:466–473. https://doi.org/10.14348/molcells.2015.0019

Choi J-W, Liao X, Stothard P, et al (2014) Whole-Genome Analyses of Korean Native and Holstein Cattle Breeds by Massively Parallel Sequencing. PLOS ONE 9:e101127. https://doi.org/10.1371/journal.pone.0101127

Danilov P, Panchenko D, Tirronen K (2020) ый льчйФкдии (The reindeer of Eastern Fennoscandia). Petrozavodsk: KarRC RAS. 187 p. [In Russian]

Edgar RC (2004) MUSCLE: multiple sequence alignment with high accuracy and high throughput. Nucleic Acids Res 32:1792–1797. https://doi.org/10.1093/nar/gkh340

Ewels P, Magnusson M, Lundin S, Käller M (2016) MultiQC: summarize analysis results for multiple tools and samples in a single report. Bioinformatics 32:3047–3048. https://doi.org/10.1093/bioinformatics/btw354

Felsenstein J (2005) PHYLIP (Phylogeny Inference Package) version 3.6. Distributed by the author. Department of Genome Sciences, University of Washington, Seattle.

Guindon S, Dufayard J-F, Lefort V, et al (2010) New Algorithms and Methods to Estimate Maximum-Likelihood Phylogenies: Assessing the Performance of PhyML 3.0. Syst Biol 59:307–321. https://doi.org/10.1093/sysbio/syq010

Härkönen S, Bisi J (2007) Suomen metsäpeurakannan hoitosuunnitelma. Maa-ja metsätalousministeriö 9/2007. [In Finnish]

Heikura K (1997) Some aspects on the recent changes in the Kuhmo-Lake Kiitehenjärvi subpopulation of the wild forest reindeer (Rangifer tarandus fennicus Lönnb.). In: Ecosystems, fauna and flora of the Finnish-Russian Nature Reserve Friendship. Finnish Environment Institute

Heino MT, Salmi A-K, Äikäs T, et al (2021) Reindeer from Sámi offering sites document the replacement of wild reindeer genetic lineages by domestic ones in Northern Finland starting from 1400 to 1600 AD. J Archaeol Sci Rep 35:102691. https://doi.org/10.1016/j.jasrep.2020.102691

Iacolina L, Corlatti L, Buzan E, et al (2019) Hybridisation in European ungulates: an overview of the current status, causes, and consequences. Mammal Rev 49:45–59. https://doi.org/10.1111/mam.12140

Klokov K (2007) Reindeer husbandry in Russia. Int J Entrep Small Bus 4:726–784. https://doi.org/10.1504/IJESB.2007.014981

Lachance J, Vernot B, Elbers CC, et al (2012) Evolutionary history and adaptation from high-coverage whole-genome sequences of diverse African hunter-gatherers. Cell 150:457–469. https://doi.org/10.1016/j.cell.2012.07.009

Lee T-H, Guo H, Wang X, et al (2014) SNPhylo: a pipeline to construct a phylogenetic tree from huge SNP data. BMC Genomics 15:162. https://doi.org/10.1186/1471-2164-15-162

Li H, Durbin R (2009) Fast and accurate short read alignment with Burrows–Wheeler transform. Bioinformatics 25:1754–1760. https://doi.org/10.1093/bioinformatics/btp324

Mallet J (2005) Hybridization as an invasion of the genome. Trends Ecol Evol 20:229–237. https://doi.org/10.1016/j.tree.2005.02.010

Montonen M (1974) Suomen Peura, 1st edn. WSOY. 118 p. [In Finnish]

Mykrä-Pohja S, Niemi M (2022) Pitkäjänteistä metsäpeuran suojelua -Metsästäjälehti. https://metsastajalehti.fi/riista/pitkajanteista-metsapeuran-suojelua/. Accessed 15 Jun 2023

Niemi M, Rautiainen M, Kilpeläinen P, Turtinen E (2021) Metsäpeuran rotupuhtaustyö ja sen kehittäminen 2017-2019. Metsähallitus, Vantaa. [In Finnish]

Paasivaara A, Hyvärinen M, Timonen P, et al (2021) Metsäpeurakanta kasvussa -Metsästäjälehti. https://metsastajalehti.fi/riista/metsapeurakanta-kasvussa/. Accessed 14 Jun 2023

Paasivaara A, Pusenius J, Ruha L, Timonen P (2022) Hirven lentolaskenta – pohjoisen erikoisuus -Metsästäjälehti. https://metsastajalehti.fi/riista/hirven-lentolaskenta-pohjoisen-erikoisuus/. Accessed 14 Jun 2023

Panchenko DV, Paasivaara AKJ, Hyvarinen MSA, Krasovskij YA (2021) The wild forest reindeer, Rangifer tarandus fennicus, in the Metsola Biosphere Reserve, Northwest Russia. Nat Conserv Res 3aпoBeдHaя Hкay 6:116–126. https://doi.org/10.24189/ncr.2021.026

Panchenko DV, Tirronen K, Danilov P, et al (2017)OцeHкaчиcл ehhocTииpacпpeдeлeHиeлecHoгo ceBepHoгo л (Rangifer taranus fennicus Lönnb.)BКлии (Assessment of the forest reindeer (Rangifer taranus fennicus Lönnb.) number and distribution in the Republic of Karelia). Becthик OxoToBeдeHия 14:156–165

Pokharel K, Weldenegodguad M, Dudeck S, et al. (2023). Whole genome sequencing provides novel insights into the evolutionary history and genetic adaptations of reindeer populations in northern Eurasia. bioRxiv

Pöllänen AT, Pakanen V-M, Paasivaara A (2023) Survival and cause-specific mortality in adult females of a northern migratory ungulate. Eur J Wildl Res 69:60. https://doi.org/10.1007/s10344-023-01686-y

Purcell S, Neale B, Todd-Brown K, et al (2007) PLINK: A Tool Set for Whole-Genome Association and Population-Based Linkage Analyses. Am J Hum Genet 81:559–575

Rambaut A (2018) FigTree v1. 4.4 (https://github.com/rambaut/figtree/releases/tag/v1.4.4)

Randi E (2005) Management of Wild Ungulate Populations in Italy: Captive-Breeding, Hybridisation and Genetic Consequences of Translocations. Vet Res Commun 29:71–75. https://doi.org/10.1007/s11259-005-0025-1

Randi E (2008) Detecting hybridization between wild species and their domesticated relatives. Mol Ecol 17:

Rankama T, Ukkonen P (2001) On the early history of the wild reindeer (Rangifer tarandus L.) in Finland. Boreas 30:131–147. https://doi.org/10.1111/j.1502-3885.2001.tb01218.x

Røed KH, Kvie KS, Bårdsen B-J, et al (2021) Historical and social–cultural processes as drivers for genetic structure in Nordic domestic reindeer. Ecol Evol 11:8910–8922. https://doi.org/10.1002/ece3.7728

Schaefer, J. A., Bergman, C. M. & Luttich, S. N. 2000. Site fidelity of female caribou at multiple spatial scales. Landscape Ecology 15:731–739. https://doi.org/10.1023/A:1008160408257

Somenzi E, Ajmone-Marsan P, Barbato M (2020) Identification of Ancestry Informative Marker (AIM) Panels to Assess Hybridisation between Feral and Domestic Sheep. Animals 10:582. https://doi.org/10.3390/ani10040582

Stothard P, Choi J-W, Basu U, et al (2011) Whole genome resequencing of black Angus and Holstein cattle for SNP and CNV discovery. BMC Genomics 12:559–559. https://doi.org/10.1186/1471-2164-12-559

Van der Auwera GA, Carneiro MO, Hartl C, et al (2013) From FastQ data to high confidence variant calls: the Genome Analysis Toolkit best practices pipeline. Curr Protoc Bioinforma Ed Board Andreas Baxevanis Al 11:11.10.1-11.10.33. https://doi.org/10.1002/0471250953.bi1110s43

van Oort H, McLellan BN, Serrouya R (2011) Fragmentation, dispersal and metapopulation function in remnant populations of endangered mountain caribou. Anim Conserv 14:215–224. https://doi.org/10.1111/j.1469-1795.2010.00423.x

Weldenegodguad M, Pokharel K, Ming Y, et al (2020) Genome sequence and comparative analysis of reindeer (Rangifer tarandus) in northern Eurasia. Sci Rep 10:8980. https://doi.org/10.1038/s41598-020-65487-y

Zheng X, Levine D, Shen J, et al (2012) A high-performance computing toolset for relatedness and principal component analysis of SNP data. Bioinforma Oxf Engl 28:3326–3328. https://doi.org/10.1093/bioinformatics/bts606

