## Supplemental tables for "Pure wild forest reindeer (*Rangifer tarandus fennicus*) or hybrids? A whole-genome sequencing approach to solve the taxonomical status"

**Supplementary Table S1** Sample information

| **Sample** | **Population** |
| --- | --- |
| FR8 | Domestic reindeer; Finland |
| FR9 | Domestic reindeer; Finland |
| FR10 | Domestic reindeer; Finland |
| FR11 | Domestic reindeer; Finland |
| FR12 | Domestic reindeer; Finland |
| FR13 | Domestic reindeer; Finland |
| FR14 | Domestic reindeer; Finland |
| FR15 | Domestic reindeer; Finland |
| FR16 | Domestic reindeer; Finland |
| FR17 | Domestic reindeer; Finland |
| RTF6 | Forest reindeer (*R. tarandus fennicus*); Finland |
| RTF7 | Forest reindeer (*R. tarandus fennicus*); Finland |
| RTF17 | Forest reindeer (*R. tarandus fennicus*); Finland |
| RTF31 | Forest reindeer (*R. tarandus fennicus*); Finland |
| RTF84 | Forest reindeer (*R. tarandus fennicus*); Finland |
| RTF96 | Forest reindeer (*R. tarandus fennicus*); Finland |
| Individual 1 | Studied *Rangifer* individual number 1 |
| Individual 2 | Studied *Rangifer* individual number 2 |
| Individual 3 | Studied *Rangifer* individual number 3 |
| Individual 4 | Studied *Rangifer* individual number 4 |
| Alaska | Alaska wild caribou |

**Supplementary Table S2** Summary of whole genome sequencing clean reads and mapping statistics

|  | **Sample** | **Read length (bp)** | **Clean reads (M)** | **Clean** | **Coverage** | **Mapping (%)** |
| --- | --- | --- | --- | --- | --- | --- |
| Finnish semi-domestic reindeer | FR8 | 150 | 265 | 39.75 |  | 98.57 |
|  | FR9 | 150 | 290 | 43.60 |  | 98.74 |
|  | FR10 | 150 | 351 | 52.72 |  | 98.13 |
|  | FR11 | 150 | 355 | 53.39 |  | 98.63 |
|  | FR12 | 150 | 299 | 45.0 |  | 98.76 |
|  | FR13 | 150 | 326 | 49.04 |  | 98.50 |
|  | FR14 | 150 | 274 | 41.13 |  | 98.66 |
|  | FR15 | 150 | 275 | 41.31 |  | 98.55 |
|  | FR16 | 150 | 254 | 38.23 |  | 98.7 |
|  | FR17 | 150 | 285 | 42.88 |  | 98.66 |
| Finnish wild forest reindeer | RTF6 | 150 | 288 | 43.26 |  | 98.5 |
|  | RTF7 | 150 | 282 | 42.40 |  | 98.63 |
|  | RTF17 | 150 | 280 | 42.01 |  | 97.83 |
|  | RTF31 | 150 | 276 | 41.55 |  | 90.71 |
|  | RTF84 | 150 | 227 | 34.07 |  | 98.74 |
|  | RTF96 | 150 | 289 | 43.47 |  | 98.43 |
| Studied individuals | Individual 1 | 150 | 239 | 35.96 |  | 98.82 |
|  | Individual 2 | 150 | 205 | 30.81 |  | 99.45 |
|  | Individual 3 | 150 | 289 | 43.40 |  | 99.28 |
|  | Individual 4 | 150 | 209 | 31.40 |  | 99.32 |
| Alaskan wild caribou | Alaska | 150 | 196 | 29.47 |  | 98..99 |

**Supplementary Table S3**. Summary of identified SNPs and Indels. TS/TV refers to transition-to-transversion mutations detected in each animal.

|  | **Sample** | **TS/TV** | **Total SNPs** | **Heterozygous** | **Homozygous** | **Indels** |
| --- | --- | --- | --- | --- | --- | --- |
| Finnish semi-domestic reindeer | FR8 | 1.89 | 8231150 | 5251861 | 2979289 | 1277079 |
|  | FR9 | 1.89 | 8340499 | 5342898 | 2997601 | 1304495 |
|  | FR10 | 1.89 | 8483154 | 5421326 | 3061828 | 1340971 |
|  | FR11 | 1.89 | 7993441 | 4990225 | 3003216 | 1225882 |
|  | FR12 | 1.89 | 7721813 | 4738619 | 2983194 | 1168592 |
|  | FR13 | 1.89 | 7947726 | 4960373 | 2987353 | 1205404 |
|  | FR14 | 1.89 | 7521587 | 4452694 | 3068893 | 1129322 |
|  | FR15 | 1.89 | 7590978 | 4564575 | 3026403 | 1134854 |
|  | FR16 | 1.89 | 7378372 | 4360595 | 3017777 | 1100895 |
|  | FR17 | 1.89 | 7665944 | 4640136 | 3025808 | 1156948 |
| Finnish wild forest reindeer | RTF6 | 1.91 | 8258004 | 4588538 | 3669466 | 1256699 |
|  | RTF7 | 1.91 | 8199400 | 4548151 | 3651249 | 1242836 |
|  | RTF17 | 1.9 | 8280182 | 4756550 | 3523632 | 1243891 |
|  | RTF31 | 1.91 | 8002516 | 4419409 | 3583107 | 1197569 |
|  | RTF84 | 1.91 | 7844002 | 4326964 | 3517038 | 1167211 |
|  | RTF96 | 1.91 | 8390203 | 4811790 | 3578413 | 1272886 |
| Studied individuals | Individual 1 | 1.91 | 8721513 | 5199135 | 3522378 | 1351760 |
|  | Individual 2 | 1.91 | 8452618 | 4966211 | 3486407 | 1289509 |
|  | Individual 3 | 1.91 | 8391229 | 4518303 | 3872926 | 1302198 |
|  | Individual 4 | 1.91 | 8338148 | 4472676 | 3865472 | 1285043 |
| Alaskan wild caribou | Alaska | 1.93 | 8433153 | 4967230 | 3465923 | 1072832 |
